# A reassessment of consensus clustering for class discovery

**DOI:** 10.1101/002642

**Authors:** Yasin Senbabaoglu, George Michailidis, Jun Z. Li

## Abstract

Consensus clustering (CC) is an unsupervised class discovery method widely used to study sample heterogeneity in high-dimensional datasets. It calculates “consensus rate” between any two samples as how frequently they are grouped together in repeated clustering runs under a certain degree of random perturbation. The pairwise consensus rates form a between-sample similarity matrix, which has been used (1) as a visual proof that clusters exist, (2) for comparing stability among clusters, and (3) for estimating the optimal number (K) of clusters. However, the sensitivity and specificity of CC have not been systemically studied. To assess its performance, we investigated the most common implementations of CC; and compared CC with other popular methods that also focus on cluster stability and estimation of K. We evaluated these methods using simulated datasets with either known structure or known absence of structure. Our results showed that (1) CC was able to divide randomly generated unimodal data into pre-specified numbers of clusters, and was able to show apparent stability of these chance partitions of known cluster-less data; (2) for data with known structure, the proportion of ambiguously clustered (PAC) pairs infers the known number of clusters more reliably than several commonly used K estimating methods; and (3) validation of the optimal K by choosing the most discriminant genes from the discovery cohort and applying them in an independent cohort often exaggerates the confidence in K due to inherent gene-gene correlations among the selected genes. While these results do not yet prove that any of the published studies using CC has generated false positive findings, they show that datasets with subtle or no structure are fully capable of producing strong evidence of consensus clustering. We therefore recommend caution is using CC in class discovery and validation.

## Author Summary

Consensus clustering (CC) is rapidly becoming the algorithm of choice for unsupervised class discovery with genomic datasets. It has been used both as a visualization tool and an inference tool, and has been cited ∼600 times since its introduction in 2003. In a typical application, The Cancer Genome Atlas (TCGA) Research Network used CC to analyze gene expression data of glioblastoma and identified four subtypes. But as often occurred in this type of studies, neither the strength of the evidence nor the sensitivity of the method was quantitatively evaluated. Here, by comparing the TCGA dataset with a series of randomly simulated datasets known to lack cluster, we highlight the potential for CC to generate false positive results in subtype discovery. We describe a CC-based summary statistic, the proportion of ambiguous clustering (PAC), as the measure to infer the optimal number of clusters. Using simulated data with known number of clusters we show that PAC outperforms commonly used methods such as CDF, Δ(K), Silhouette Width, GAP-PC and CLEST in scenarios closely resembling real studies. We conclude by making practical recommendations for conducting unsupervised class discovery using CC.

## Introduction

Cluster analysis is one of the main tools for unsupervised subtype discovery from high-dimensional data. Since 1996, cluster analysis of microarray-derived gene expression profiles has led to the recognition of molecular subtypes of many cancers [1–6]. However, it is difficult to quantify clustering strength by casting it into a standard hypothesis-testing problem, because each real dataset could have a unique covariance structure. For example, shared regulatory pathways inevitably produce strong gene-gene correlations, thus a multivariate Gaussian distribution without gene-gene correlation does not constitute a valid null in practice. This difficulty led to the development of non-parametric, resampling-based methods, where multiple subsamples of the original dataset are clustered, and the results are compared to assess cluster stability. One such method, CLEST [7], computes cross-validation errors for a range of potential cluster numbers (denoted K throughout this study), and compares these errors to find the optimal K. Another resampling-based method, consensus clustering (CC) [8], has recently gained widespread application in subtype discovery. CC calculates a “consensus rate” between all pairs of samples, defined as the frequency with which a given pair is grouped together in multiple clustering runs under a certain degree of random permutation, implemented either by random initialization or by sample-or gene-subsampling. The resulting similarity matrix is often used as both a visualization tool and an inference tool for putative clusters, where the differences between within-group and between-group consensus rates allow the assessment of cluster stability and the estimation of the optimal K.

The main assumption of CC is that if the objects under study were drawn from distinct sub-populations that truly exist, different subsamples of these objects would exhibit similar cluster numbers, i.e., the true K. This assumption is easily validated in cases of well-separated clusters. However, whether apparently robust clusters might also arise from structureless data (i.e., having no clusters) has not been studied. This question underscores a potential limitation of resampling-based methods, namely the difficulty of formally evaluating the significance of cluster results without stating the underlying null assumptions. Although this limitation is acknowledged in the literature [for example, 8], many studies still relied on the consensus rate heatmap to visually demonstrate the existence of clusters, without investigating the robustness of the conclusions.

## A motivating example

Glioblastoma multiforme (GBM) was the first cancer type studied by The Cancer Genome Atlas (TCGA) Research Network [9], and was reported to have four molecular subtypes according to gene expression clusters discovered by CC [10]. Here we use the same data in a technical reassessment of CC. We will not discuss biological implications of GBM subtypes, which have been revised and expanded since the initial study by TCGA [11–13].

We ran CC on the gene expression data for the first GBM cohort (n = 202, referred to as GBM1, see **Materials and Methods**), with K = 4 and k-means as the base clustering method. The consensus rate matrix was calculated by 500 repeated clustering runs, taking a random 80% subset of genes (Figure 1a) or samples (Figure 1b) in each run. As originally reported [10], the consensus heatmaps (Figure 1a-b) show four crisp clusters; and it was the crispness of the clusters that was cited as the strong evidence for inherent structure at K = 4 in GBM1. However, the appearance of clusters in the Pearson’s correlation coefficient matrix (Figure 1c) is substantially weaker, with many samples having strong correlations with samples in a different cluster. Similarly, principal component analysis (PCA) (Figure 1d) does not show distinct gaps among the four reported clusters, rather they represent contiguous partitions of an unbroken data cloud. These findings raise the question whether CC has over-stated the robustness of clusters.

**Figure 1.**
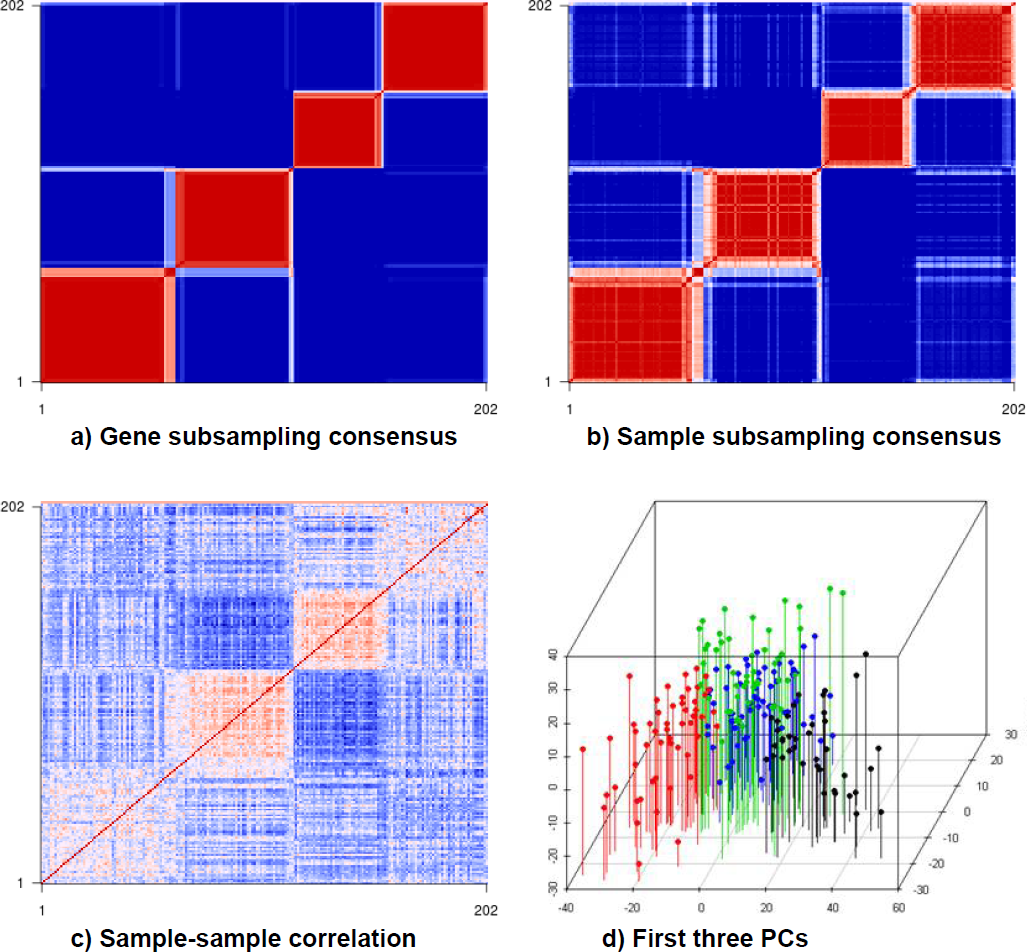
Different ways to visualize the clustering signal in GBM1. (a) gene-subsampling consensus heatmap with K = 4, (b) sample-subsampling consensus heatmap with K = 4, (c) sample-sample correlation heatmap, (d) four k-means clusters visualized along PC1-PC2-PC3 (x-axis, z-axis and y-axis). The variances explained by PC1-PC2-PC3 are 21.6%, 9.9%, and 7.9% respectively. The color scale on consensus heatmaps ranges from 0 to 1, where 0 corresponds to blue, 1 corresponds to red, and 0.5 corresponds to white. The same color scale is used throughout the paper unless otherwise stated.

A related issue is the robustness of estimated K. We re-ran CC using K = 2 and K = 3, applied average-linkage hierarchical clustering to the consensus matrices, and re-displayed the same correlation matrix in Figure 1c with the hierarchical-clustering-based order for K = 2 (Figure 2a), K = 3 (Figure 2b) and with the order from the first principal component scores (Figure 2c). Each panel in Figure 2 showed interesting structure, suggesting that it is possible to re-order a sample-sample similarity matrix in different ways to support different claims regarding optimal K.

**Figure 2.**
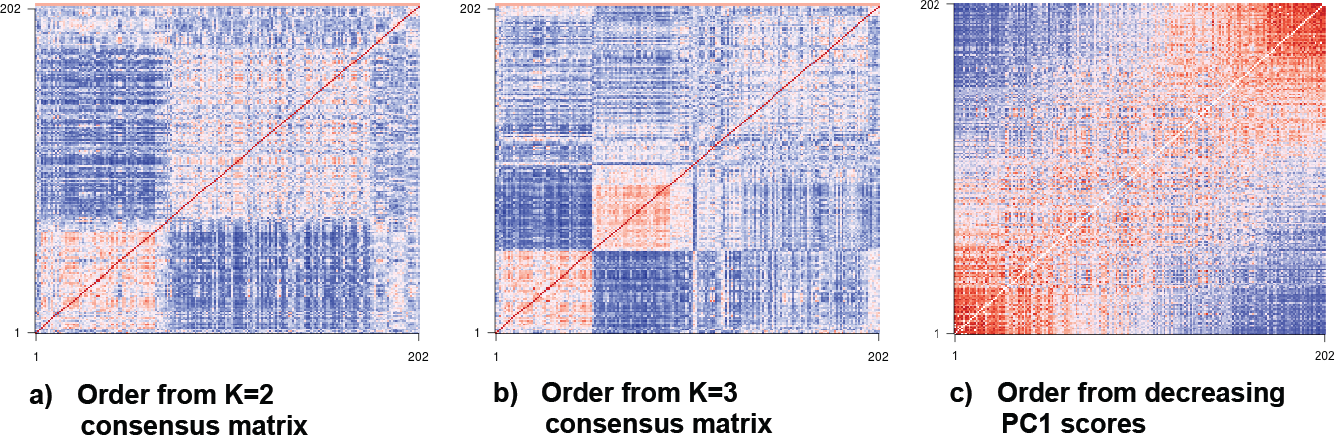
GBM1 sample-sample correlation matrix ordered three different ways. The order of samples is obtained from (a) average-linkage HCLUST on the K = 2 gene subsampling consensus matrix (b) average linkage HCLUST on the K = 3 gene subsampling consensus matrix (c) decreasing PC1 scores. It is easy to re-order the same sample-sample matrix in different ways to support different hypotheses about structuredness.

The examples above motivated us to ask the following questions: (1) How can an investigator know if he/she is merely partitioning data from a unimodal distribution into multiple groups? (2) How should the optimal K be determined? (3) How to verify the existence of clusters and how to validate K? In the following, we address these questions by systematically studying the sensitivity and specificity of CC on known negative and known positive datasets.

## Results

### CC is capable of finding clusters in null datasets of *unimodal* distribution

We consider the typical case where the input data are expression levels of p genes measured on n samples, and the question is whether the n samples form K clusters. The underlying assumption of CC is that repeated subsampling of genes or samples can capture the true structure of the population from which the data come from. However, since CC does not generate a measure of statistical confidence associated with this estimation procedure, one needs to compare clustering results for the test dataset with those from an ensemble of negative datasets, which form a null distribution. Several types of null distribution have been proposed in the literature, but they vary in how closely they mimic real-life data values and gene-gene correlations. For example, the null n-p matrix can be populated randomly from a univariate **uniform** or **unimodal** distribution [7]. Alternatively, if multivariate distributions are used, the covariance among the variables can be either set to zero or “borrowed” from real datasets, by sampling random projections in the principal component space derived from the real data. In the following we will examine how the choice of the null model could affect the reported confidence of clustering results, beginning with an illustrative example.

We tested the performance of CC on two simple datasets, (1) 100 samples that form a regularly spaced square-shaped grid in the PC1-PC2 space (called *Square1*), and (2) ∼300 samples forming a similar but circle-shaped grid (called *Circle1*). Briefly, we drew two 1000-element random vectors from Normal(0,1) that served as fixed PC1 and PC2 vectors. Next, for *Square1*, we generated 100 pairs of [PC1, PC2] coefficients that would place 100 samples onto a 10-by-10 grid in the PC1-PC2 space. In this formation, samples had regularly increasing PC1 scores from left to right in the PC1-PC2 plot, and regularly increasing PC2 scores from bottom to top. The [PC1, PC2] scores were slightly “wiggled” from the grid points by adding random Normal(0,1) noise. The final 100 × 1000 matrix is formed by linear combinations of the two fixed PC vectors with the 100 different coefficient pairs. Similarly, for *Circle1*, we repeated the procedure above but changed the number of samples from 100 to 400, forming a 20-by-20 grid plus the same level of random wiggle. We then trimmed the square grid to keep only the samples with a distance to the center smaller than a radius of ∼9.62 grid units, leaving ∼300 samples that form a circle. Strictly speaking, both *Square1* and *Circle1* have higher gene-gene correlations than a matrix filled with Normal(0,1) data because all 100 (or 300) objects are derived from the same PC1 and PC2 vectors. However, they are still unimodal in the sense that the sample placements lack local compactness or separation. Thus they can serve as the test case where no cluster is known to exist, and from which no robust cluster should be found.

We performed consensus clustering on *Circle1* for K = 2-8, using k-means as the base method. In Figure 3, the upper panels show the group partition in a single typical k-means run; and lower panels show the CC matrix heatmaps for 250 runs with 80% subsampling. While there is no inherent structure in *Circle1*, CC can nonetheless partition the samples into K subgroups, which are spatially well segregated. Importantly, CC is able to show a high apparent stability of clusters in the heatmap, for example, at K = 2-4 (Figure 3a-3c). Moreover, the stability is further improved for larger K (such as 7 or 8), making it tempting to conclude that the original data contain 7 or 8 clusters (Figure 3f-3g).

**Figure 3.**
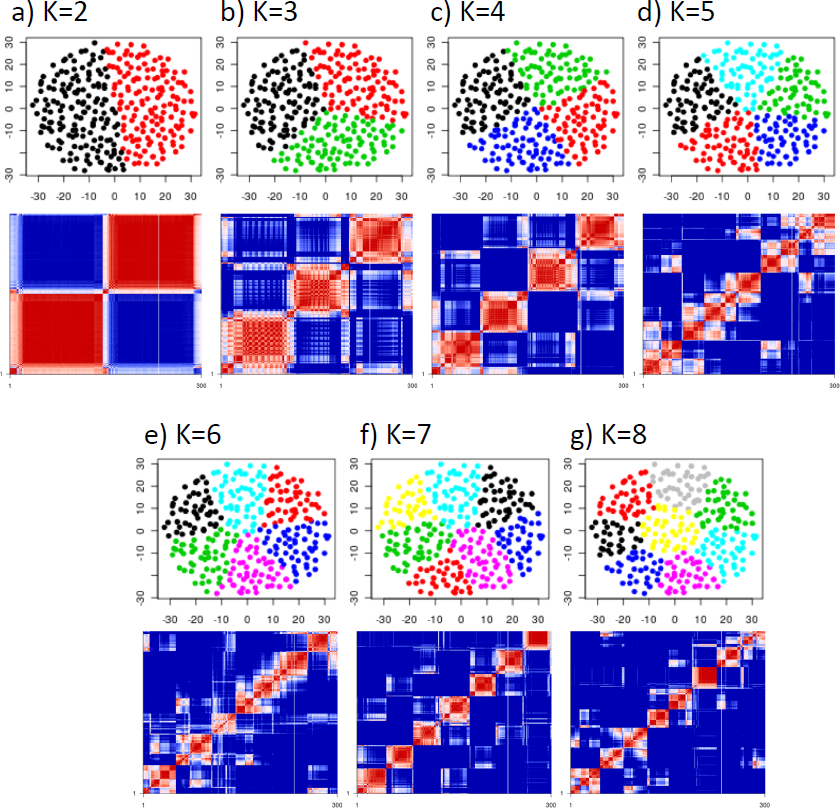
Consensus heatmaps can show apparent clusters even in unimodal distributions. **Top** panels in (a-g) show Circle1 k-means partitioning for K = 2-8 displayed on PC1 (17.7%) on the x-axis vs PC2 (15.1%) on the y-axis. **Bottom** panels show consensus heatmaps for K = 2-8 with 80% sample subsampling and k-means as the base method. Visual evidence alone can be misleading; hence formal approaches are needed to test validity.

The apparent stability in this example is potentially caused in part by the presence of outliers or “corners” of the sample distribution. To explicitly investigate this, we performed CC on *Square1* for K = 2-5. As in *Circle1*, *Square1* samples show clear partitioning and apparent stability, especially for K = 4 (Figure 4). Clusters for K = 2-3 were not as ‘clean’, suggesting that the four corners of the grid helped to anchor the K = 4 partitions and lend them stability.

**Figure 4.**
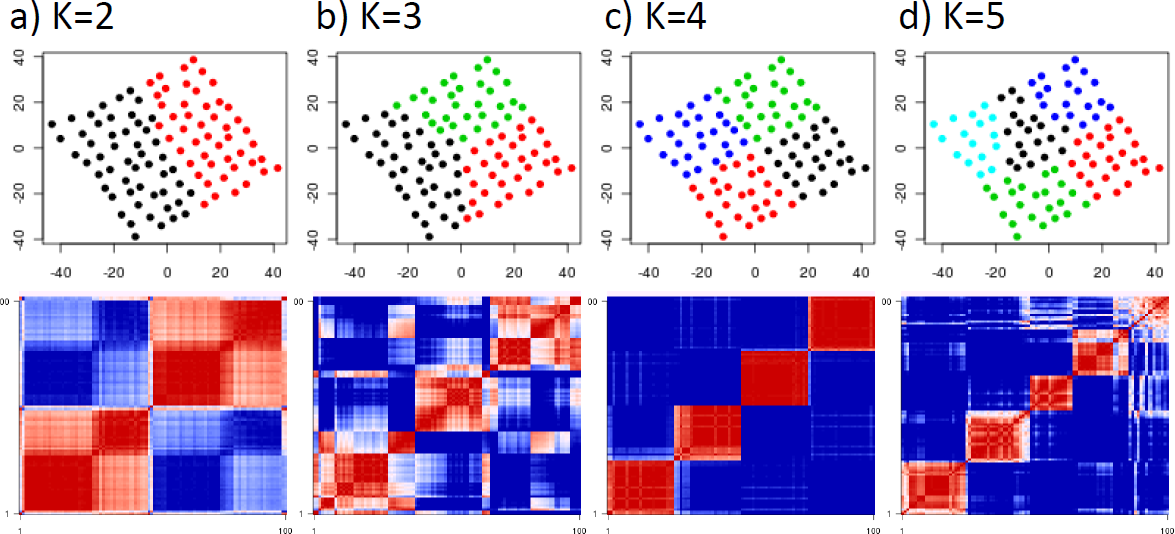
Consensus heatmaps show apparent clusters for certain K values in the unimodal Square1 distribution. **Top** panels in (a-d) show Square1 k-means partitioning for K = 2-5 displayed on PC1 (21.8%) on the x-axis vs PC2 (19.1%) on the y-axis. **Bottom** panels in (a-d) show consensus heatmaps for K = 2-5 with 80% sample subsampling and k-means as the base method. In (c), the four corners of the data cloud act as anchors and make the clusters look more stable on consensus heatmaps.

Together, these simple illustrative examples show that CC is able to claim apparent stability of chance partitioning of null datasets drawn from a unimodal distribution, and thus has the potential to lead to over-interpretation of cluster stability in a real study. A related lesson is that, visual evidence alone can be misleading, and formal inference methods are needed to test the robustness of the clusters. This is particularly relevant in practice, as many published studies utilizing CC neglected to evaluate the strength of the evidence, and relied on visualization of the CC matrix to declare clusters. In the next section, we construct a more realistic null model by applying GBM1’s gene-gene correlation structure to generate a family of datasets obeying a unimodal distribution. We then evaluate the original GBM1 data in comparison with these empirical null datasets.

### CC shows stable clusters for null models harboring empirical gene-gene correlations

In a data matrix with p genes in one dimension and n samples in another, gene-gene correlation (a p-p matrix), and sample-sample correlation (an n-n matrix) are dependent of each other. For example, if the samples fall into two clusters, the genes that differentiate the two clusters will be correlated, leading to a recognizable structure in gene-gene correlation. Conversely, if a group of genes are co-regulated, they will limit the “shape” of sample projections in the p-dimensional space. For example, if gene-1 (g1) and gene-2 (g2) are strongly correlated, samples will tend to occupy an elongated ellipsoid in the g1-g2 dimension rather than a sphere, making it easier to identify sample clusters from one end of the ellipsoid to another. In short, gene-gene correlation is a key parameter in the unsupervised discovery of sample clusters, and needs to be considered when constructing realistic null distributions.

In settings naturally encountered in genomic studies, n << p, and gene-gene correlation information is often reliably represented by the top eigenvectors, i.e., the principle component *loadings* that quantify the contribution of each of the p genes to the most salient data structure. When the top K eigenvectors are known from a real dataset, one can create null cluster-less datasets with the same gene-gene correlation by (1) drawing the top K principal component scores of n simulated samples by randomly sampling a univariate Gaussian distribution to form an n-K score matrix, and (2) multiplying this score matrix with the K eigenvectors from the real dataset (**Materials and Methods**). By repeating this procedure, we generated 50 null datasets from GBM1 and called this collection the **pcNormal** datasets. When needing to run one-to-one comparisons with GBM1, we chose a representative dataset from pcNormal as the one for which the silhouette width statistics (defined in **Materials and Methods**) is ranked 25th among the 50, and denoted this dataset as **Sim25**.

Although the pcNormal datasets lack structure, CC analysis show stable clusters with K = 2, 3, 4. An example, Sim25 (Figure 5) showed “block-like” clusters in the K = 4 heatmap, and these are as crisp as those for the original GBM1 data (compare Figure 5b-5c with Figure 1a-1b). Although this comparison does not establish that GBM1 has no structure, it shows that simulated data with no known local density or outlier groups are fully capable of producing visually convincing evidence for strong clusters via the use of CC.

**Figure 5.**
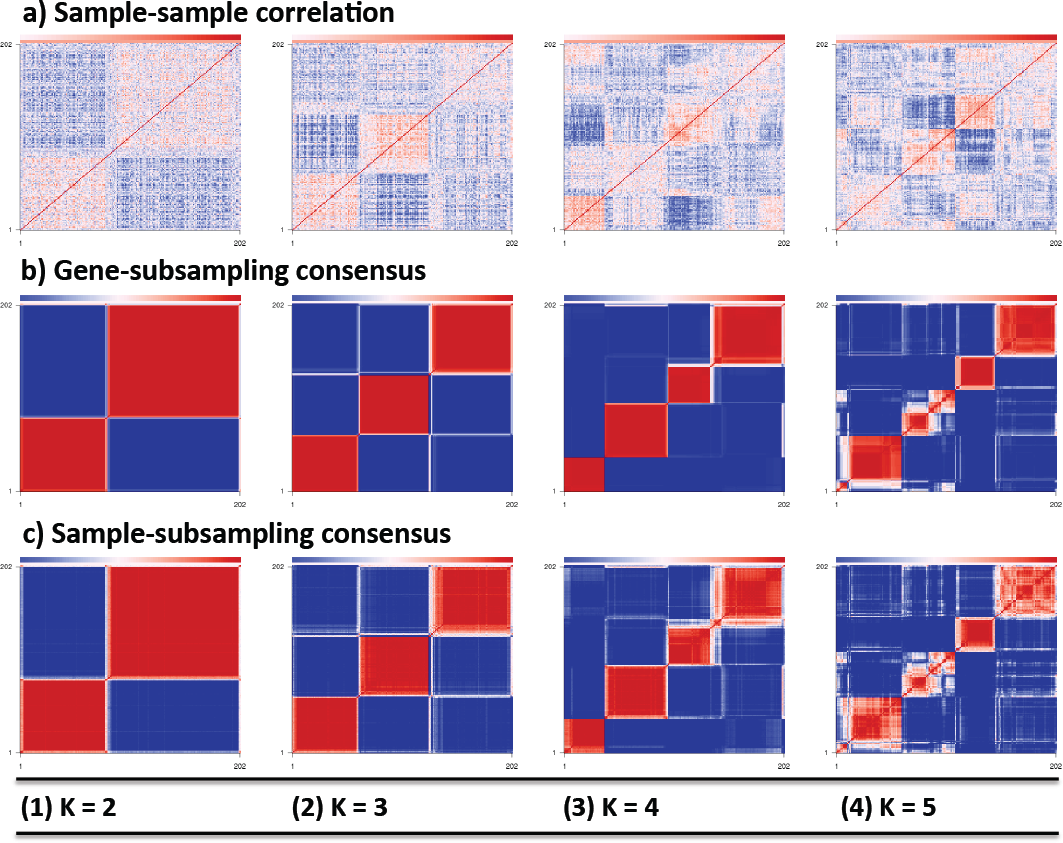
CC shows stable clusters in Sim25 even though Sim25 lacks structure. (a1-a4) sample-sample correlation heatmaps, (b1-b4) 80% gene-subsampling consensus heatmaps, (c1-c4) 80% sample-subsampling consensus heatmaps. Sim25 is chosen as a characteristic random dataset from the pcNormal null distribution. For each K in 2-5, the order of samples on all three heatmaps is the one obtained from average-linkage hierarchical clustering on the gene-subsampling consensus matrix of the relevant K value. The base clustering method for CC is k-means.

### Quantitative comparisons of cluster strength between GBM1 and the null datasets

CC heatmaps in Figure 1 and 5 allow visual comparisons of cluster strength. However, formal inference requires quantitative summaries for the presence of clusters under a range of possible Ks. Two such summaries are CLEST [7] and silhouette width [14]. Briefly, CLEST is a resampling-based method that randomly partitions the original dataset into a learning set and a test set. The former is used in an unsupervised clustering method to build a K-cluster classifier, which is applied to partition the latter (the test set) in supervised assignment. The test set is also partitioned independently using the same unsupervised clustering algorithm as applied for the training set. The concordance between the supervised and unsupervised partitions for the test set is summarized by measures such as the Fowlkes-Mallows (FM) index, for which a higher value indicates stronger clustering signals in the original data. Silhouette width is computed for each sample and each K based on the comparison of its distance to its own cluster and that to other clusters. A dataset with strong clusters tend to show a high average value of positive silhouette width, and fewer samples of negative silhouette width.

We apply these two methods to compare clustering strength between three TCGA datasets and the 50 pcNormal null datasets. The three datasets are: TCGA’s first and second GBM cohort (**GBM1** and **GBM2,** respectively), and the **validation** dataset [10], which is a combination of data from multiple prior studies [15–18]. CLEST results (Figure 6a) show that, for all K values except 2, the three real datasets have higher FM values than the null datasets. GBM1, in particular, show the highest FM values, suggesting that GBM1 has more structure than the null datasets. However, K = 4 is not clearly the optimal number of clusters for GBM1, because the differences from the null datasets are comparable across K = 3 − 8.

**Figure 6.**
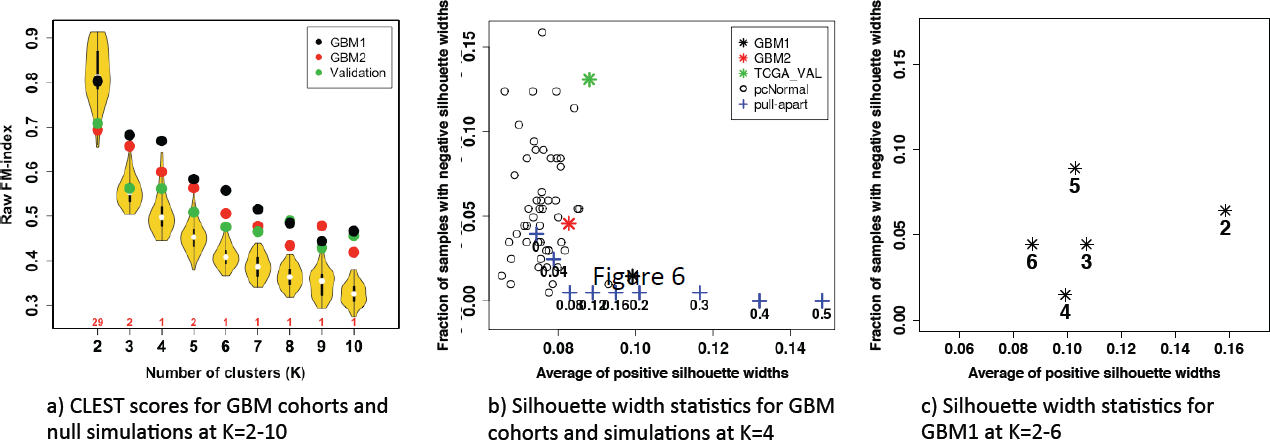
CLEST and silhouette width analysis. (a) CLEST results for GBM1, GBM2, Validation, and pcNormal null datasets in the range K = 2-10. (b) K = 4 silhouette width analysis for GBM1, GBM2, Validation, pcNormal null datasets, and pull-apart positive datasets. (c) Silhouette width analysis for GBM1 in the range K = 2-6. In (a), GBM1’s FM-score from CLEST beats those of pcNormal simulations at as early as K = 3. In (b) the x-axis shows the average of positive silhouette widths, and the y-axis shows the fraction of negative silhouette widths. GBM1 is within the range of 50 pcNormal simulations (shown with hollow circles) along the y-axis. However, it appears as an outlier along the x-axis when compared with these null datasets. The pull-apart degree for positive datasets ranges from 0 to 0.5 (shown with blue plus symbols). Along the x-axis, GBM1 is close to the positive dataset with pull-apart degree 0.2. In (c), it is not possible to declare K = 4 as optimal.

Figure 6b shows the average of positive silhouette widths on the x-axis and the fraction of samples with negative silhouette widths on the y-axis. Datasets with strong clustering signals are expected to appear on the lower-right side of the plot. At K = 4, GBM1 is within the distribution of the 50 pcNormal datasets along the y-axis, but is a positive outlier along the x-axis. We also added to this figure the results from simulated **positive** datasets with four known clusters pulled apart with controlled degrees of separation (as measured by a, explained below and in **Materials and Methods**). GBM1 is most similar to the dataset with a = 0.2. This result suggests that GBM1 has a certain clustering structure, however it does not confirm K = 4 as the optimal number of cluster in the range K = 2-6 using silhouette width statistics (Figure 6c). GBM2 and the Validation dataset are within the range of pcNormal for both axes, reinforcing the results from Figure 6a that they have weaker structures than GBM1.

The apparent structure of GBM1 can be attributed to the fact that our simulations relied on normally distributed samples in the hyperspace and could not capture all the spatial features of the actual dataset. For example, Figure 1d showed “protrusions” of GBM1 samples towards two lower front corners of the PC1-PC2-PC3 cubic space. Such local formations in the hyperspace are difficult to match by simulations, partly explaining the observed difference between GBM1 and the null simulations (Figure 6b). In short, while some quantitative measures, such as CLEST and average silhouette width, could bring out the uniqueness of GBM1, other measures, such as the heatmaps in Figure 1a-1b and the negative silhouette width fraction, could not. This underlines the fact that different clustering measures emphasize different features of a given heterogeneous high-dimensional dataset. The average of positive silhouette widths, for example, is strongly influenced by the existence of one or more highly compact clusters.

### Limitations of Δ(K), GAP-PC and CLEST for finding K

CC matrix has been used as a sensitive heuristic for visualizing clusters, assessing stability, and inferring the optimal K. To formally estimate K, two statistics derived from CC, the cumulative distribution function (CDF) and the proportional change in the area under the CDF curve upon an increase of K (Δ(K)) have been proposed (explained in more detail in **Materials and Methods**). We investigated the performance of CDF and Δ(K), along with two other methods, using simulated positive datasets of known clustering separation and known K.

To generate datasets of known structure, we first obtained K clusters in Sim25 using a k-means run, and then gradually “pulled apart” the samples in each cluster, in PC space, from the global center of all samples. We implemented this pulling apart procedure for K = 2-6, with pull-apart degree “a” in the range [0, 0.8], where 1.0 represents pulling the PC scores of the samples away from the global mean by a distance equal to the original distance between the cluster mean and the global mean.

We tested four methods for estimating K: CDF, Δ(K), GAP-PC [19], and CLEST, shown in four different rows in Figure 7. Within each row, arranged from left to right are the results from four datasets, for no-pull-apart at a = 0, 2-way pull-apart at a = 0.08, 3-way pull-apart at a = 0.12, and 4-way pull-apart at a = 0.12, respectively. These a values were chosen as the smallest values in the range [0,0.8] where the CDF plot exhibits a flat curve for the true K value (Figure 7a).

**Figure 7.**
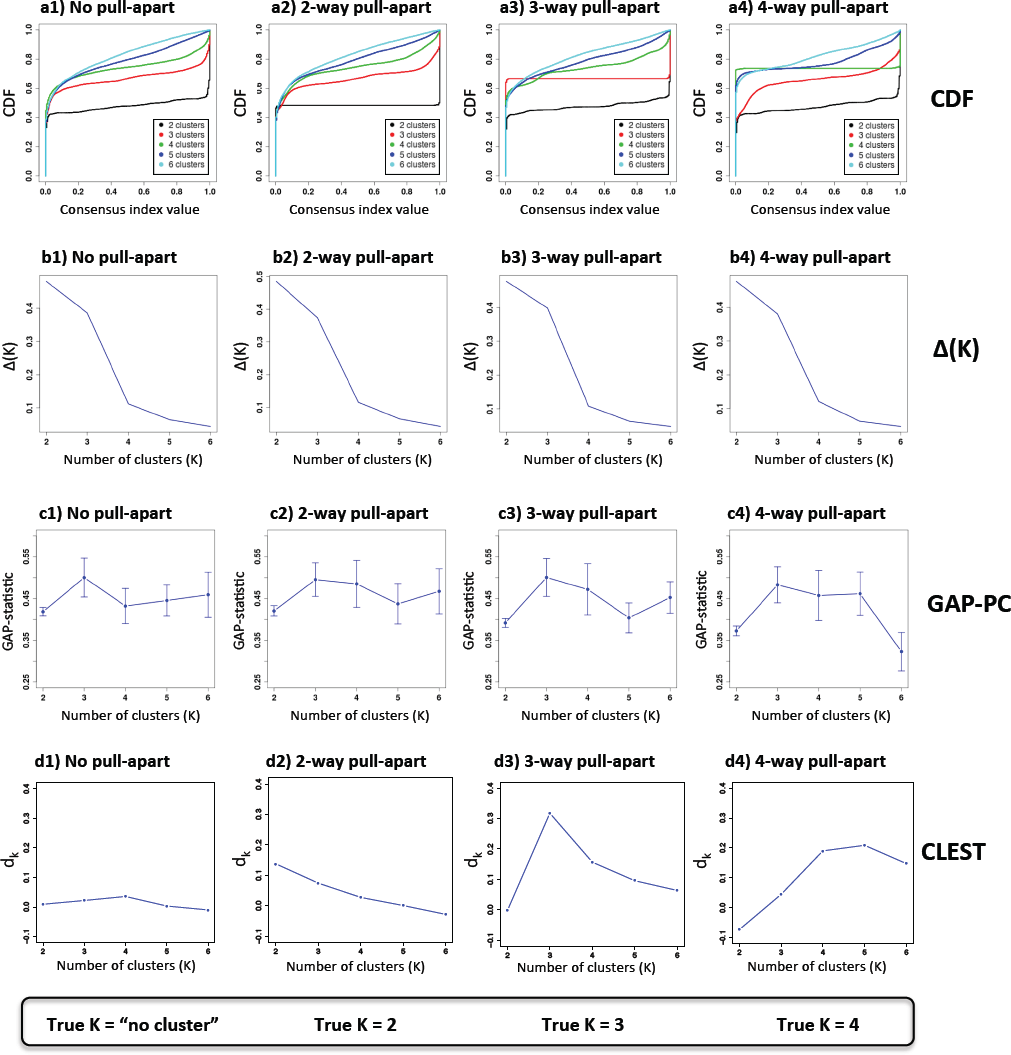
Δ(K) plots are inadequate for revealing the optimal K. The four columns in this dataset, from left to right, belong to (1) a randomly generated unimodal dataset, (2) a 2-way pull-apart dataset with degree of pull-apart = 0.08, (3) a 3-way pull-apart dataset with degree of pull-apart = 0.12, and (4) a 4-way pull-apart dataset with degree of pull-apart = 0.12. **First row (a1-a4):** CDF plots from the consensus matrices. CDF curves for K = 2-6 are shown with black, red, green, blue and cyan lines respectively. For the pull-apart datasets in (a2,a3,a4), the CDF curve for true K shows perfect 0s and 1s while the unimodal dataset in (a1) does not have such a curve. **Second row (b1-b4):** Δ(K) plots across K = 2-6. An elbow occurs at K = 4 in all of these plots suggesting an optimal K value of 3. **Third row (c1-c4):** GAP plots across K = 2-6. In all four figures, the optimal K value according to the original interpretation is 3. **Fourth row (d1-d4):** CLEST plots across K = 2-6. *d*^*k*^ (y-axis) is the difference between the observed similarity statistic and its estimated expected value under the null hypothesis of K = 1. The decision criterion involving *d*^*k*^ suggest an optimal K of 1, 2, 3, and 5 in these four pull-apart cases respectively.

As shown in Figure 7a, CDF is able to infer the correct number of clusters, as indicated by the flat CDF curves of only the true K values (for K = 2 − 4), reflecting a perfect or near-perfect partitioning of the dataset at the correct K. As expected, the no-pull-apart dataset in 7a does not have such a flat curve because it is constructed with K = 1.

In contrast, all Δ(K) curves in Figure 7b are alike in that they all exhibit an ‘elbow’ at K = 4, suggesting that K = 4 had smaller improvement than K = 3, and that K = 3 could be called optimal, even when the true K is 1, 2, or 4.

The GAP-statistic provides an estimate of K by comparing the change in within-cluster dispersion with that expected under a reference null distribution. There are two alternative algorithms: for *GAP-unif*, the null datasets are generated from a uniform distribution over the range of each observed feature; for *GAP-PC*, they are generated from a uniform distribution in the principal component space [19]. The authors suggested in [19] that the optimal K is the smallest K for which the GAP score is larger than the lower bound for K + 1; where the lower bound is defined as the GAP score minus the standard error for that particular K value. According to this inference rule, all four plots in Figure 7c conclude an optimal K of 3, even when the true K is 1, 2, or 4.

The CLEST method is based on the *d*_*k*_ statistic, which measures the difference between *t*_*k*_, the observed similarity statistic (such as the FM index), and *t*_*k*_^0^, its estimated expected value under the null hypothesis of K = 1. Among the K values that satisfy a pre-specified *d*_*min*_ criterion (here *d*_*min*_=0.05), the optimal K is the one with maximum *d*_*k*_. If none of the tested K values satisfy the pre-specified criteria, the optimal K is concluded to be 1. In figure 7-d1, optimal K is 1 because *d*_*k*_ < 0.05 for all K. For the 2, 3, and 4-way pull-apart cases, CLEST concludes an optimal K of 2, 3, and 5 respectively, as given by the K with the maximum *d*_*k*_. In total, CLEST was able to make correct inferences in three out of four cases tested.

In summary, when clusters are sufficiently separated, the CDF curves exhibit a flat middle segment only for the true K, and this can be used to infer the optimal K (Figure 7a). In contrast, Δ(K) can be uninformative even in the presence of genuine structure (Figure 7b). The published GAP decision criterion may also perform poorly (Figure 7c). CLEST, on the other hand, may have similar sensitivity compared with CDF curves (Figure 7d). In a later section we will expand our evaluation to a wider range of (K, a) combinations.

### Proportion of ambiguous clustering (PAC) and its performance

In the CDF curve of a consensus matrix, the lower left portion represents sample pairs rarely clustered together, the upper right portion represents those almost always clustered together, whereas the middle segment represent those with ambiguous assignments in different clustering runs. As shown in Figure 7a, the CDF curves show a flat middle segment only for the true K, suggesting that very few sample pairs are ambiguous when K is correctly inferred. To quantify this feature of the CDF curve we developed the “proportion of ambiguous clustering” (PAC), defined as the fraction of sample pairs with consensus indices falling in the intermediate sub-interval (u_1_, u_2_) ∈ [0, 1]. A low value of PAC indicates a flat middle segment, and a low rate of discordant assignments across permuted clustering runs. We therefore can infer the optimal K by finding the lowest PAC.

Figure 7 showed results for four particular combinations of K and a. To compare different methods across a wider range of (K, a) values, we developed a new plot, showing five panels of stacked bar plots for each of six methods (Figure 8a-f). For each method, the five panels correspond to, from bottom to top, K = [2,…,6]. Within each panel, from left to right are segmented bar plots for increasing a in the range [0, 0.08]. Within each bar plot, the length of the vertical segments show the fraction of inferred K across 50 simulated datasets for the given (K, a) combination. The segments were color-coded to facilitate direct visualization of how well the inferred Ks agree with the true K, as shown on the far right. Such plots allow systematic performance comparisons for different methods under different (K, a). Figure 8 shows the results for PAC, Δ(K), CLEST, GAP-PC with the original decision rule, GAP-PC with a modified decision rule (explained in **Materials and Methods**), and the silhouette width. For a given (K, a), the same 50 datasets were used in testing the six methods.

**Figure 8.**
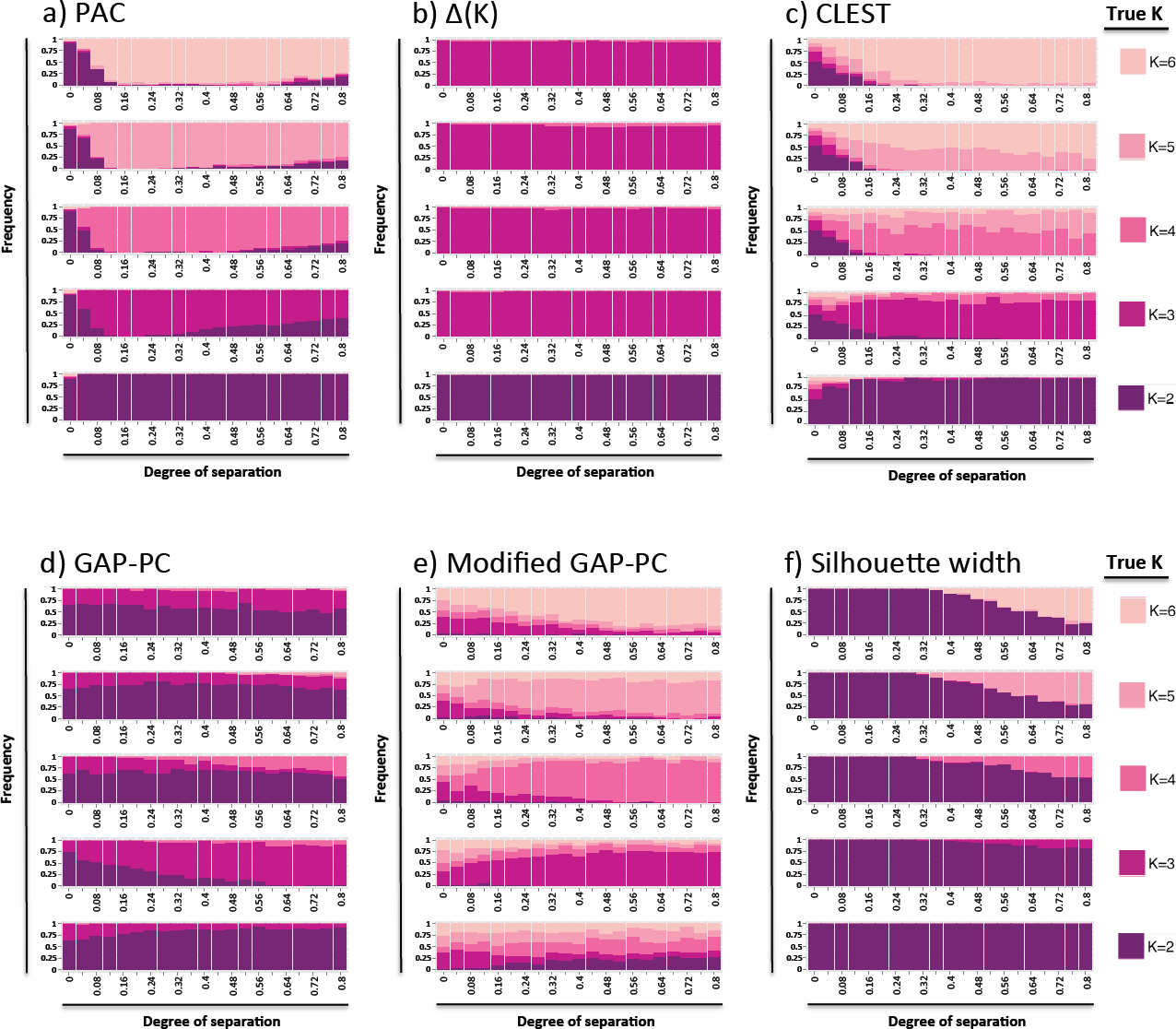
The identifiability zone for PAC is drastically better than other measures/methods. Identifiability graphs for (a) PAC (probability of ambiguous clustering), (b) Δ(K), (c) CLEST, (d) GAP-PC with the original decision rule, (e) GAP-PC with a modified decision rule, and (f) silhouette width. The x-axis shows the strength of the real pull-apart signal, denoted with **a**. The y-axis shows the true number of pull-apart clusters in the dataset, denoted with K. The colors in the bars indicate estimated K values for the corresponding (K,a) pair. The length of each color in a given bar is proportional to the frequency of that particular K value in the set of 50 simulations.

As shown in Figure 8a, PAC performs well across most of tested (K, a) pairs. When a > 0.04 it detects the correct K for K = 2-6. When a < 0.04 it tends to call K = 2 even for larger true Ks, and this tendency to under-call returned for some of the runs for larger a.

In comparison, Δ(K) detects the correct K for K = 2 and 3, but calls K = 3 even when the true K = 4−6 (Figure 8b), i.e., it consistently under-calls when K >3. This result is consistent with that in Figure 7b.

For CLEST, the inferred K is correct for most datasets with true K = 2,3,6 and with a > 0.2 (Figure 8c). When a < 0.2 it calls K = 2 even for true K of 3-6. When the true K is 4, CLEST has a tendency to overcall K = 5. The tendency to overcall is even stronger when the true K is 5, in which case CLEST is more likely to call K = 6 than K = 5. On the whole, the parameter space of correct identification is smaller than in PAC, but much bigger than other methods.

The original GAP-PC method performs well for K= 2-3, and improves with larger a, but it severely under-calls for K = 4−6 (Figure 8d). In contrast, the modified GAP-PC performs well for K = 3−6, although it tends to over-call when true K = 2 (Figure 8e). On the whole, the modified GAP-PC is much improved over the original GAP-PC and ranks second best (after PAC) among the six methods.

Lastly, the silhouette width severely under-calls in most situations. For a > 0.4, however, it calls either K = 2 or the correct K, never calling a third value of K (Figure 8f).

In sum, using simulated data with known number of clusters we show that PAC outperforms several commonly used methods in calling the correct K.

### Gene-gene correlation among most discriminant genes makes it easy to “validate” any K

After an optimal K is determined for a dataset, the next task is to validate K. This can be difficult when there is no external information (e.g., known class labels) with which to calculate classification error rates. An alternative solution is to replicate the observation of K clusters in independent datasets. Ideally, the replication in the test set should not “borrow” any information from the first, learning set. However, a method that has become highly popular involves (1) determining the most discriminant genes from the original dataset for its optimal K value, and (2) using these same genes to classify samples in an independent dataset. In a popular implementation [2,17], the best classifier genes for each of K clusters are chosen from the learning set, and a heatmap of all learning samples with only these genes is constructed, with the samples and the genes both grouped in K clusters. Next, another heatmap is made using the same genes for the replication samples. Observing the same number of discrete gene and sample clusters in the latter heatmap is considered a validation of K. We show below that, due to the gene-gene correlation structure in genomic datasets, this approach can easily “validate” a K value even for data with no known structure.

For this analysis, we chose Sim25, the representative dataset from the pcNormal simulations, as the “original” dataset from which the clusters and discriminating genes were to be learned. Following the procedure in [10], we first run k-means on Sim25 with K = 4 and obtain four clusters for the 202 samples. We then find the 210 most discriminating genes for each cluster based on the t-scores for each cluster against the three other clusters. The four gene sets are combined to form a list of 551 unique genes, and are used in both Sim25 and the replication datasets. The heatmap of Sim25 (Figure 9a) shows discrete placement of four gene sets and four sample classes. However, for nine null datasets from the pcNormal simulations, chosen in a way to represent the entire spectrum of silhouette widths, the same clustering signature is observed in all 9 cases (Figure 9b). This is because the most discriminant genes contain many that are strongly correlated with each other. Such correlations arise from co-regulation by common upstream regulators, or from inherent differences in different cell types, and can easily recur in an independent dataset even when the clustering pattern is different or absent. Results in Figure 9 show that the blocks of genes co-expression in subsets of samples could persist even when the independent dataset is simulated from a unimodal distribution, thus apparently validating K.

**Figure 9.**
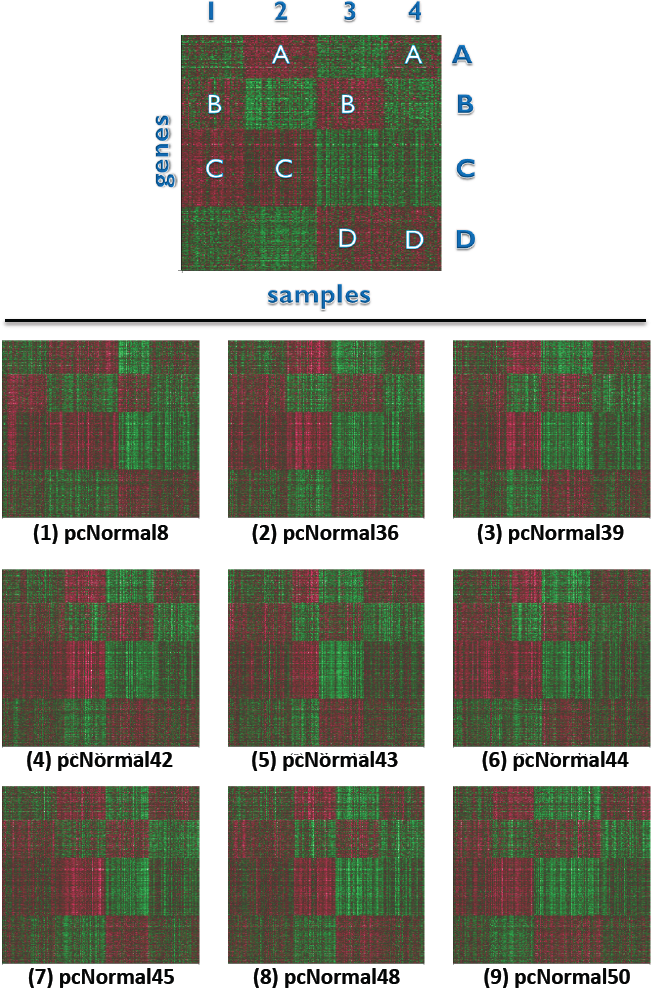
The gene signature from Sim25 is preserved even when the new data are random. (a) The heatmap of most discriminant genes (groups A-D) for k-means clustering of Sim25 with K = 4. (b) Heatmaps for nine datasets similarly simulated as Sim25. The x-axis shows samples as partitioned into 4 clusters with k-means, and the y-axis shows the same ‘most discriminant genes’ from Sim25. These nine datasets, although they are null, were able to show the same visual placement for the gene signature blocks in Sim25.

## Discussions

CC measures cluster reproducibility under perturbed runs, and provides a meaningful heuristic for visualizing cluster stability. However, it is important to distinguish the utility of CC as a formal inference tool from its informal, visualization function. Our assessment using simulated Circle1 and Square1 has shown that CC is exquisitely sensitive: declaring structure where there is no significant separation or local compactness. This led us to systemically study the strength of clustering by comparing the real data with suitably formed null datasets.

We compared GBM1 with random unimodal data and observed that the consensus clustering heatmaps and summary statistics for GBM1 were often within the empirical null distribution. Methods such as CLEST and average silhouette width were able to distinguish GBM1 from the null datasets, likely because of the local clusters or outliers in GBM1. However, methods such as Δ(K), GAP-PC and negative silhouette fraction either could not confirm the structure in GBM1 or could not confirm optimal K = 4. These findings are not surprising, as different clustering measures and validation procedures emphasize different features of the data. For instance, the average-silhouette-width is strongly influenced by the existence of one or more highly compact clusters. The GAP statistic, on the other hand, is affected by the degree of overlap between clusters. Thus it is important to be aware of the strengths and weaknesses of each method, especially when clusters have poor separation. Moreover, we found that, while the consensus matrix by itself is not a suitable inference tool, one of its distribution features, the proportion of ambiguously clustered (PAC) pairs, reflected the true structure of the data better than other common strategies such as CDF, Δ(K), silhouette width, GAP-PC, and CLEST.

Another novelty of this assessment is to evaluate different methods across a range of “positives”, i.e., data with known structures, with varying degrees of separation (a) and number of clusters (K). However, to limit the scope of our analysis we had to make some specific assumptions: (1) Samples in cluster boundaries are assigned to a single cluster; no partial memberships are used, (2) clusters are viewed as co-equal, without any hierarchy, and (3) clusters are simulated with similar sizes, with no outliers added. These complicating factors need to be explored in future studies.

A choice of null distribution depends on the two distinct tasks of class discovery: first, to determine if there is evidence for structure; second, when it is shown that substructure does exist, to determine the optimal number of clusters. For the first task, a ***global null*** should be constructed to test the “structure vs. no-structure” hypotheses, and needs to account for the gene-gene correlation structure in the original dataset as it affects the shape of the sample distribution in the high-dimensional space, potentially driving the apparent cluster stability. Here we refrain from using the terms “random” and “homogeneous” to describe this type of global null, because the gene-gene correlations can be considered as a form of innate data structure. For the second task, a set of ***study-specific null*** datasets for alternative K’s should be used, because K cannot be reported as optimal unless the null hypotheses of K − 1 and K + 1 are both rejected.

In summary, CC can be a powerful tool for identifying clusters, but it needs to be applied with caution as it is prone to over-interpretation. If clusters are not well separated, CC could lead one to conclude apparent structure when there is none, or declare cluster stability when it is subtle. To reduce false positive in the exploratory phase of a new study, we recommend the following:

- Do not rely solely on the consensus matrix heatmap to declare the existence of clusters, or to estimate optimal K.
- Do a formal test of cluster strength using simulated unimodal data with the same gene-gene correlation as in the empirical data.
- Apply the proportion of ambiguous clustering (PAC) as a simple yet powerful method to infer optimal K.
- Do not use the most discriminant genes for K clusters in the test dataset to validate K in a new dataset.

## Materials and Methods

### Datasets

This study covered three cohorts of GBM samples. GBM1 is the cohort analyzed by the TCGA pilot study [9,10]. Gene expression data were downloaded from http://tcga-data.nci.nih.gov/docs/publications/gbm_exp/. Most of our analyses were based on “unifiedScaledFiltered.txt”, which contains processed data for 1740 most variable genes for 202 GBM samples. A second cohort was subsequently analyzed by TCGA and was called GBM2. Gene expression data for GBM2 were downloaded from the TCGA Data Matrix webpage (https://tcga-data.nci.nih.gov/tcga/dataAccessMatrix.htm). This dataset contains 175 samples, and we focused on the same 1740 genes as in GBM1. The third cohort was the validation dataset used in [10] and is a collection of samples from four previous studies [15–18]. This dataset, called “validation” in this work, contains 260 samples, and the number of genes in common between GBM1 and **validation** is 1676. This dataset was also downloaded from http://tcga-data.nci.nih.gov/docs/publications/gbm_exp/.

### Generating null distributions based on empirical gene-gene correlations

By using PCA we decomposed the GBM1 data of 202 samples and 1740 genes into (1) the 202 × 202 principal component score matrix and (2) the 202 × 1740 eigenvector matrix. When simulating new datasets, in order to preserve the relative magnitude of the PC scores for different PCs in GBM1, we constructed 202 × 202 random PC score matrices by populating each column with random draws from a univariate Gaussian distribution with mean = 0 and standard deviation equal to that of the corresponding column in the original GBM1’s score matrix. Multiplying this random score matrix with the 202 × 1740 eigenvector matrix yields a null 202 × 1740 dataset in which it is known that no cluster exists. We repeated this procedure 50 times to generate a null collection of *pcNormal* datasets. Each dataset in the collection forms an ellipsoid in the 1740-dimensional space.

The steps for this procedure are as follows:

1. Using principal component analysis, we obtain the orthogonal matrix *A* of GBM1 eigenvectors.

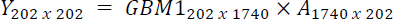 *Y* is the PC score matrix for GBM1. A is the PC vector matrix.
2. Next, we simulate a random score matrix *Y*^*N*^ where column *i* is distributed normally with zero mean and standard deviation equal to that of column *i* in Y.

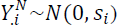 where *s*_*i*_ is the standard deviation of *y*_*i*_ and *i* = {1, …, 202}.
3. Multiplying *Y*^*N*^ with the transpose of *A* yields *Q*^*N*^, one of the *pcNormal* simulations.

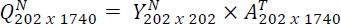
4. We repeat steps 2 and 3 50 times to obtain a collection of 50 *pcNormal* simulations.

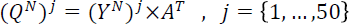

### Choosing a representative null dataset from pcNormal

A representative dataset, called **Sim25,** is chosen from *pcNormal* as having clustering signals closest to the median of the 50 datasets, as measured by the average of positive silhouette widths and the fraction of negative silhouette widths. Let

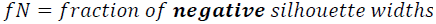

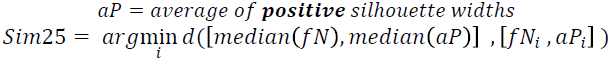

where [*fN*_*i*_, *aP*_*i*_] is the silhouette width statistics for simulation *i* ∈ {1, …,50} and *d* is the Euclidean distance function in the [*fN*, *aP*] space.

### Choosing nine pcNormal simulations for validation by most discriminant genes

The Euclidean distance of the (aP,fN) pair to the median of these quantities in the cohort was computed for each of the 50 simulations and ranked from lowest to highest. Every 5th dataset was selected among the ranked simulations, such that the [6, 11, 16, 21, 26, 31, 36, 41, 46] datasets were chosen. This ensures that nine datasets cover the entire range of clustering strength in *pcNormal*.

### Generating positive datasets for performance comparison

To generate a positive dataset with K clusters, we first ran k-means on Sim25 with the desired K value. Then, we computed the centroids of the PC scores for each of the K clusters, and added a known fraction of the centroid coordinates (i.e. the pull-apart degree, denoted as a positive scalar, “a”) to the original PC scores of samples in the corresponding cluster. Next, we multiplied the resulting PC scores from all clusters by the original principal component vectors of Sim25 so that the pull-apart datasets preserve the initial gene-gene correlation structure (with the caveat that increasing a values would gradually increase the gene-gene correlation).

Algorithmically, we execute the following steps for this procedure:

1. Use principal component analysis to obtain the eigenvector matrix *A* as before.

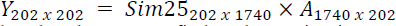
2. Use a partitioning method such as k-means to find K clusters in Sim25, assign each sample *s*_*i*_ (*i* = 1, …,202) into one of K classes. The set of samples in class *k* (*k* = 1, …, *k*) is denoted as *E*_*k*_.
3. For each class *E*_*k*_, compute the centroid *C*_*k*_ of PC scores 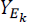.
4. For each class *E*_*k*_ and for a given pull-apart degree *a*, compute pulled-apart score matrix 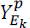.

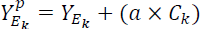
5. Multiply *Y*^*p*^ with *A*^*T*^ to obtain the pulled-apart dataset *X*^*p*^.

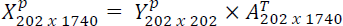

### Base methods for consensus clustering: K-means

Given a set of observations (***x***_**1**_, ***x***_**2**_, …, ***x***_***n***_) where each observation is a d-dimensional real vector, k-means clustering aims to partition the *n* observations into *k* sets (*k* ≤ *n*), ***S*** = {*S*_1_, *S*_2_, …, *S_k_*} so as to minimize the within-cluster dispersion:

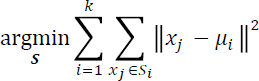

where *μ*_*i*_ is the mean of points in *S*_*i*_ [20].

The method starts with k arbitrary cluster centers. Each step consists of labeling data points with their nearest cluster center, and updating the centers of the new clusters. The procedure stops when the clusters formed at two consecutive steps are the same.

## Five ways to measure clustering signals

### Empirical CDF

For a given consensus matrix *M*, the corresponding empirical cumulative distribution (CDF) can be defined over the range [0, 1] as follows:

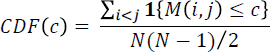

where **1**{…} denotes the indicator function, *M(i, j)* denotes entry (i, j) of the consensus matrix *M*, *N* is the number of rows (and columns) of *M*, and *c* is the consensus index value [8].

### Proportional area change under CDF (Δ(K))

The changes of CDF as K increases provide evidence for finding the optimal number of clusters. A CDF curve that closely describes a three-phase step function as mentioned before is indicative of a higher cluster stability. A method for using this information is to select the largest K that induces a large enough increase in the area under the CDF [8] which is defined as:

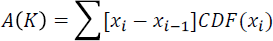

The progression, in turn, can be visualized by plotting the proportion increase Δ(*K*) in the CDF area as K increases. Δ(*K*) is computed as follows:

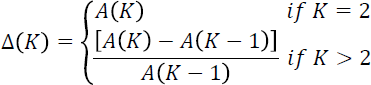

### Silhouette width

The silhouette widths of a clustering result [14] have been applied to report clustering strength and to find the optimal number of clusters K. For an object i in the dataset, let A denote the cluster to which it is assigned, and define

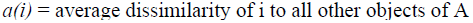

For each of the clusters *C* ≠ *A*, calculate

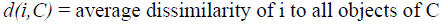

Then select the smallest of d.

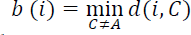

The silhouette width of object i is defined as:

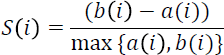

It can be seen that S(i) lies between −1 and + 1.

We chose to compare GBM1 with the null simulations according to two summary statistics derived from silhouette widths. One is the “*fraction of samples with negative silhouette widths*”. A negative silhouette width indicates that the sample is likely to have been assigned to the wrong cluster. The second statistic is the “*average of positive silhouette widths*”. Higher values of this statistic indicate stronger cluster separation.

### GAP-statistic

The GAP-statistic provides an estimate for the number of clusters in a dataset by comparing the within-cluster dispersion *W*_*k*_ with that expected under an appropriate reference null distribution (*W*_*k*_^*b*^ where *b* ∈ {1,2, …, *B*}) [19]. We first computed *W*_*k*_ for each number of clusters *K* ≥ 2. We have not included K = 1 to ensure comparability across all methods tested here; methods such as CDF and silhouette width do not allow an inference of K = 1.

For the reference distribution, there are two alternative algorithms: *GAP-unif* and *GAP-PC*. For the former, the null datasets are generated from a uniform distribution over the range of each observed feature; and for the latter, they are generated from a uniform distribution over a box aligned with the principal components of the centered design matrix. The first approach has the advantage of simplicity, but the second can factor in the shape of the data distribution [19].

In this study, we generated *B* = 40 reference datasets according to the *GAP-PC* algorithm as it can take into account the shape of the data distribution. Next, we computed the within-cluster sum of squares *W*_*k*_^1^, …, *W*_*k*_^*B*^ for each reference dataset and estimated the *gap*_*k*_ statistic with the formula:

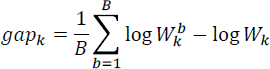

The standard error for this quantity, *S*_*k*_, was then computed as 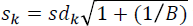 where *sd*_*k*_ is the uncorrected sample standard deviation of the log *W*_*k*_^*b*^ quantities with *b* ∈ {1,2, …, *B*}.

Finally, we chose the number of clusters via:

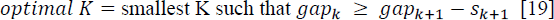

**Modified GAP-PC**: The original GAP-PC decision rule for the optimal K is to choose the smallest K where the *gap*_*k*_ score is larger than the lower bound for K + 1. A more intuitive decision rule is to declare the K value with the highest *gap*_*k*_ score as the optimal K.

### CLEST

CLEST [7] is a resampling-based method that randomly partitions the original dataset into a learning set and a test set. The former is used to build a K-cluster classifier, which is applied to partition the latter (the test set) in supervised assignment (such as DLDA [21]). The test set is also partitioned independently with an unsupervised clustering algorithm (such as k-means). The concordance between the supervised and unsupervised partitions is summarized by measures such as the Fowlkes-Mallows (FM) index, for which a higher value indicates a stronger agreement of clustering results. CLEST computes concordance scores for a range of K, and compares these to those obtained from null simulations to determine the optimal K.

